# Complementary roles of human protamine 1 and spermine in giant DNA compaction

**DOI:** 10.64898/2026.09.06.749741

**Authors:** Takashi Nishio, Yoshio Kato

## Abstract

During sperm maturation, protamines progressively replace histones and compact the paternal genome. Spermine (SPM), a naturally occurring tetravalent polyamine that induces DNA compaction, is also abundant in seminal fluid. However, how SPM and protamines jointly influence higher-order structural transitions of giant DNA remains poorly understood. Here, we investigated the cooperative effects of human protamine 1 (PRM1) and SPM using single-molecule fluorescence microscopy. PRM1 induced a continuous conformational transition from coil to globule through intermediate conformations, whereas SPM induced a discrete coil–globule transition. Notably, under coexistence conditions with low concentrations of PRM1, SPM promoted structural progression along the PRM1-induced continuous folding pathway toward the fully compact globule state. Quantitative analysis supported these observations and demonstrated distinct yet cooperative modes of DNA compaction by PRM1 and SPM. These findings demonstrate complementary roles of human PRM1 and SPM in giant DNA compaction and provide a physicochemical framework for understanding genome compaction during sperm maturation.

## 1. Introduction

During spermiogenesis, histones are progressively replaced by protamines, a family of small, highly basic DNA-binding proteins conserved among vertebrates, resulting in the extreme compaction of the paternal genome in mature sperm [1, 2]. Owing to their essential roles in sperm chromatin organization and fertility, protamine-mediated genome compaction has been extensively investigated using diverse biochemical, biophysical, and structural approaches [3-6]. DNA condensation by naturally occurring polyamines has also been extensively investigated, providing fundamental insights into the physicochemical principles governing higher-order structural transitions of DNA [7, 8]. Polyamines are ubiquitous small polycationic molecules present in virtually all living organisms and regulate diverse cellular functions through electrostatic interactions with negatively charged biomolecules, including nucleic acids [9]. Among naturally occurring polyamines, spermine (SPM), a ubiquitous tetravalent polyamine found in humans and many other organisms, is abundant in seminal fluid [10]. Nuclear SPM is also present in human spermatogenic cells and progressively decreases during spermatid differentiation, concomitant with histone-to-protamine replacement and chromatin compaction [11]. Its ability to induce higher-order structural transitions of giant DNA has been extensively characterized by single-molecule observations [8, 12, 13]. However, how SPM and protamines jointly influence higher-order structural transitions of DNA remains poorly understood.

Due to the characteristic properties of giant DNA molecules that are not readily manifested in short DNA fragments, in vitro single-molecule observations of giant DNA provide a powerful approach for directly visualizing higher-order structural transitions [8, 14]. Using this approach, Makita et al. demonstrated that salmon protamine induces a continuous conformational transition through intermediate conformations, in marked contrast to the discrete coil–globule transition induced by polyamines [15]. These observations indicate that the mode of these higher-order structural transitions depends strongly on the physicochemical properties of the condensing agents. Single-molecule studies further demonstrated that higher-order structural transitions induced by polyamines are markedly altered under coexistence conditions of biologically relevant cations, revealing competitive and cooperative effects among different cationic species [16-18]. Together, these studies establish single-molecule analysis of giant DNA as a powerful framework for investigating the cooperative regulation of higher-order structural transitions by biologically relevant DNA-condensing molecules.

In the present study, we investigated the combined effects of human protamine 1 (PRM1) and SPM on the higher-order structure of giant DNA using single-molecule fluorescence microscopy. We demonstrate that human PRM1 and SPM play distinct yet complementary roles in giant DNA compaction, with SPM promoting structural progression along the PRM1-induced folding pathway.

## 2. Materials and methods

### 2.1. Preparation of recombinant human protamine 1 (PRM1)

A synthetic DNA fragment encoding human protamine 1 (PRM1) was cloned into a modified pET expression vector containing an N-terminal polyhistidine tag, mEmerald, and bdSUMO sequence, as described previously [19]. The recombinant plasmid was transformed into *Escherichia coli* BL21(DE3). Overnight cultures were diluted 100-fold into LB broth (Lennox; Merck, Darmstadt, Germany) supplemented with 10 μg/mL kanamycin and cultured at 37 °C until the optical density at 600 nm reached approximately 0.7. Protein expression was induced by the addition of 0.1 mM isopropyl β-D-1-thiogalactopyranoside (IPTG), followed by incubation overnight at 18 °C.

Cells were harvested by centrifugation and lysed in lysis buffer (50 mM Tris-HCl (pH 8.0), 500 mM NaCl, 10% glycerol, 10 mM imidazole, 1 mM DTT, and 1 mM AEBSF) using a Digital Sonifier 250D Advanced homogenizer (Branson, CT, USA). After centrifugation, the clarified lysate was applied to a Ni-NTA affinity column (QIAGEN, Venlo, Netherlands), and the His-mEmerald-bdSUMO-PRM1 fusion protein was eluted with lysis buffer containing 500 mM imidazole. The buffer was exchanged to cleavage buffer (10 mM Tris-HCl (pH 8.0), 100 mM NaCl) using a PD-10 desalting column (Cytiva, Marlborough, MA, USA). The N-terminal fusion tag was removed by incubation with 0.2 μM recombinant His-tagged SENP1 protease overnight at 4 °C [20].

Following tag cleavage, the NaCl concentration was adjusted to 500 mM, and the reaction mixture was fractionated using an Amicon Ultra 30 kDa centrifugal filter (Merck Millipore, Burlington, MA, USA). The flow-through fraction containing PRM1 was collected and subsequently concentrated and desalted using an Amicon Ultra 3 kDa centrifugal filter by repeated dilution with 10 mM Tris-HCl (pH 8.0) until the NaCl concentration was reduced to below 1 mM. The concentration of purified PRM1 was estimated by comparing its band intensity with that of bovine serum albumin standards on SDS-PAGE. The purified PRM1 was stored at −80 °C until use. The amino acid sequence of recombinant PRM1 and representative SDS-PAGE analysis used for protein quantification are provided in Supplementary Fig. S1.

### 2.2. Single-molecule fluorescence microscopy and image analysis

T4 GT7 DNA (166 kbp; Nippon Gene, Tokyo, Japan) was prepared at 0.1 µM in nucleotide units in 10 mM Tris-HCl (pH 7.5; Nippon Gene) containing 4% (v/v) 2-mercaptoethanol (2-ME; FUJIFILM Wako Pure Chemical Corporation, Osaka, Japan). Purified PRM1 and spermine tetrahydrochloride (SPM; Nacalai Tesque, Kyoto, Japan) were added to the DNA solution at the indicated final concentrations. DNA was fluorescently stained with 0.05 µM YOYO-1 iodide (Invitrogen/Thermo Fisher Scientific, Waltham, MA, USA), following our previously reported procedure [13, 17]. Individual DNA molecules were observed using an inverted fluorescence microscope (IX71, Olympus, Tokyo, Japan) equipped with a 100× oil-immersion objective and a 1.6× intermediate magnification. Excitation light from a mercury lamp (USH-103OL, Olympus) was introduced through a NIBA filter set (Olympus; excitation, BP470–490 nm; dichroic mirror, DM505; emission, BP510–550 nm). Fluorescence images were acquired at 33.3 frames per second with an sCMOS camera (ORCA-Fusion BT, Hamamatsu Photonics, Shizuoka, Japan).

Image analysis was performed using ImageJ (National Institutes of Health, Bethesda, MD, USA). Because individual DNA molecules continuously underwent translational and conformational Brownian motion, each molecule was measured using a frame in which it was clearly focused and sufficiently separated from surrounding molecules. The long-axis length, *L*, was defined as the maximum distance across the two-dimensional fluorescence image and represents the apparent molecular extension projected onto the imaging plane. For quantitative comparison, the ensemble-averaged long-axis length, <*L*>, was calculated for each conformational state under each experimental condition. DNA conformations were classified as coil, partial globule, or globule using criteria based on fluorescence morphology and *L*, adapted from our previous single-molecule analysis [17]. Compact molecules with *L* ≤ 1.0 µm and no clearly extended segment were classified as globules. Molecules with *L* > 1.0 µm that contained one or more locally compact, high-intensity regions together with an extended segment within the same DNA molecule were classified as partial globules. Molecules with an extended fluorescence distribution but without such localized compact regions were classified as coils. Representative fluorescence images and corresponding quasi-3D intensity profiles of these conformations are shown in Fig. 1A. Overlapping molecules and exceptionally bright or large objects that were likely to represent multimolecular aggregates were excluded from the analysis. Fifty individual DNA molecules were analyzed for each condition.

**Fig. 1.**
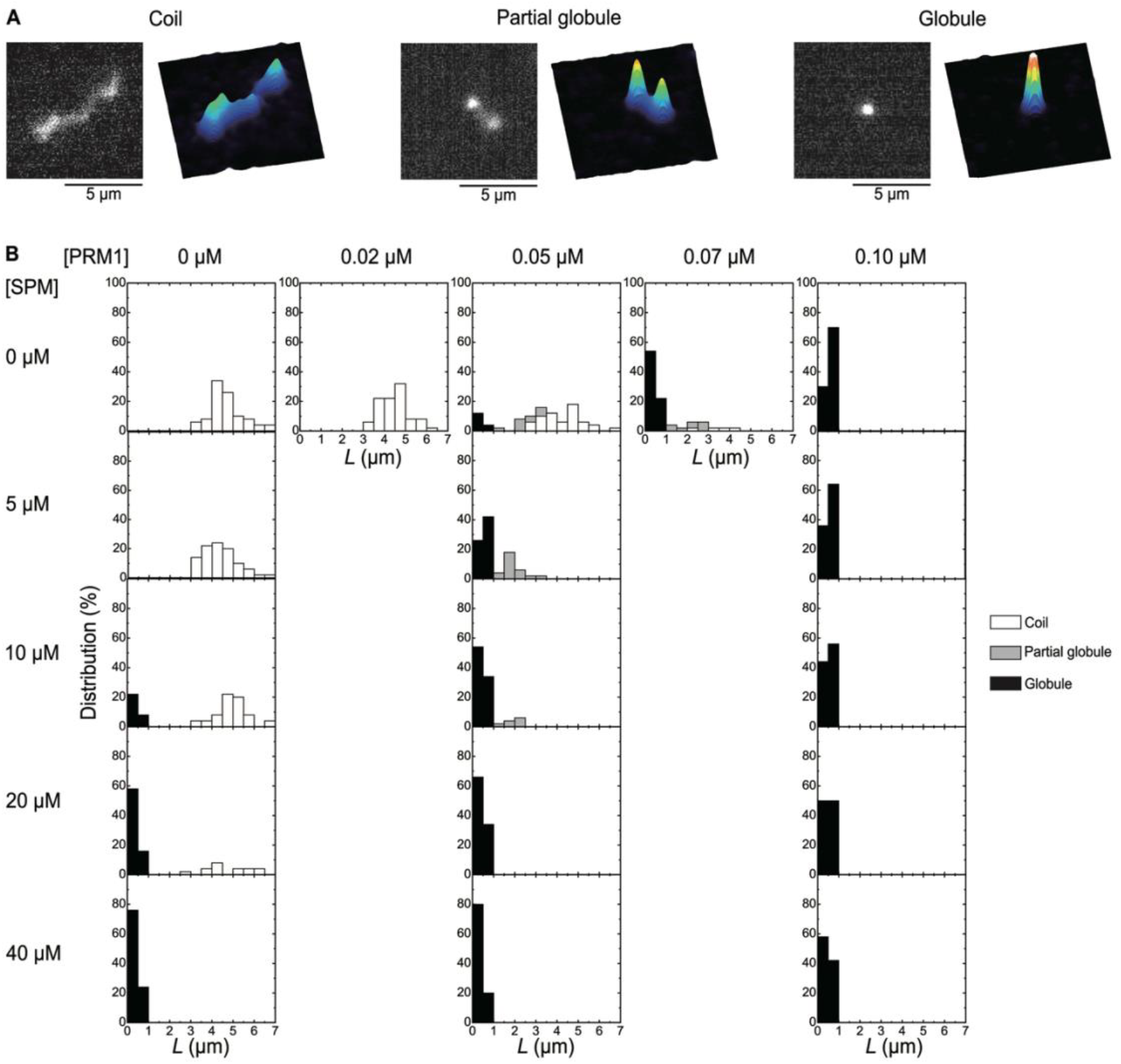
Higher-order structural transition of T4 GT7 DNA induced by PRM1 and SPM. (A) Representative fluorescence microscopy (FM) images and corresponding quasi-3D fluorescence intensity profiles of T4 GT7 DNA molecules exhibiting coil, partial globule, and globule conformations. (B) Distributions of the long-axis length, *L*, of individual DNA molecules under various PRM1 and SPM concentrations. For each condition, 50 individual DNA molecules were analyzed.

## 3. Results

### 3.1. Continuous DNA compaction induced by PRM1

Figure 1 summarizes the higher-order structural changes of individual T4 GT7 DNA (166 kbp) molecules induced by PRM1 and SPM. Representative fluorescence microscopy (FM) images together with the corresponding quasi-three-dimensional fluorescence intensity profiles of representative conformational states are shown in Fig. 1A, and the distributions of the long-axis length, *L*, under different PRM1 and SPM concentrations are presented in Fig. 1B. In the absence of PRM1, DNA molecules exhibited the coil conformation. As the PRM1 concentration increased, the conformational population was continuously redistributed from coil toward partial globule and globule states. At 0.10 μM PRM1, only globule conformations were observed. These observations demonstrate that PRM1 induces a continuous conformational transition from coil to globule through intermediate conformations. The present observations are in good agreement with previous single-molecule studies using salmon protamine, which also demonstrated a continuous conformational transition through intermediate conformations [15]. Continuous conformational changes accompanied by intermediate conformations have also been reported for giant DNA interacting with other macromolecular polycations, including histone H1 and amino-functionalized polymers [21, 22]. Together, these observations suggest that continuous transitions through intermediate conformations are a common feature of giant DNA interacting with macromolecular polycations. Recent AFM and single-molecule tethering studies further demonstrated that protamine-mediated DNA folding proceeds through multiple intermediate structures, including partial loops, loops, flowers, and loop-stack conformations [6, 23, 24]. These microscopic observations are consistent with the present fluorescence-microscopy results showing a gradual redistribution of the global DNA conformation through intermediate states. Together, these findings indicate that protamine-mediated DNA compaction proceeds through multiple intermediate conformations rather than through the discrete coil–globule transition characteristic of small multivalent polyamines [12, 13, 15].

### 3.2. Cooperative effects of PRM1 and SPM on giant DNA compaction

Based on the characteristic conformational changes induced by PRM1 alone (Fig. 1B), the effects of SPM were examined at three representative PRM1 concentrations (0, 0.05, and 0.10 μM). In the absence of PRM1, increasing the SPM concentration induced a concentration-dependent discrete coil–globule transition, and only globule conformations were observed at 40 μM SPM. This discrete transition is consistent with previous single-molecule observations of giant DNA condensed by multivalent polyamines [12, 13, 15]. In contrast, in the presence of 0.05 μM PRM1, where all three conformational states coexisted, the coil conformation was no longer observed following the addition of 5 μM SPM, and partial globule and globule conformations coexisted. Further increasing the SPM concentration increased the globule population, with a corresponding decrease in the partial globule population, and only globule conformations were observed at 20 μM SPM, whereas 40 μM SPM was required in the absence of PRM1. At 0.10 μM PRM1, all analyzed DNA molecules remained in the globule conformation throughout the examined SPM concentration range.

To further characterize the structural changes induced by PRM1 and SPM, the ensemble-averaged long-axis length, <*L*>, was analyzed separately for the coil, partial globule, and globule conformations under varying PRM1 concentrations (Fig. 2A) and SPM concentrations (Fig. 2B). In the absence of SPM, increasing the PRM1 concentration progressively decreased <*L*> of the coil conformation. The partial globule conformation also showed a tendency toward shorter <*L*> values with increasing PRM1 concentration, whereas <*L*> of the globule conformation remained nearly constant over the examined PRM1 concentration range (Fig. 2A). In contrast, SPM alone had little effect on <*L*> of either the coil or globule conformations (Fig. 2B). In the presence of 0.05 μM PRM1, <*L*> of the partial globule conformation showed a slight decrease with increasing SPM concentration, whereas <*L*> of the globule conformation again remained nearly constant. At 0.10 μM PRM1, only the globule conformation was observed, and its <*L*> remained nearly constant throughout the examined SPM concentration range (Fig. 2B). These quantitative analyses further support the distinct roles of PRM1 and SPM in DNA compaction. PRM1 progressively decreased <*L*> of the coil and partial globule conformations during the continuous folding process, whereas SPM primarily promoted a discrete coil–globule transition without appreciably altering <*L*> within each conformational state.

**Fig. 2.**
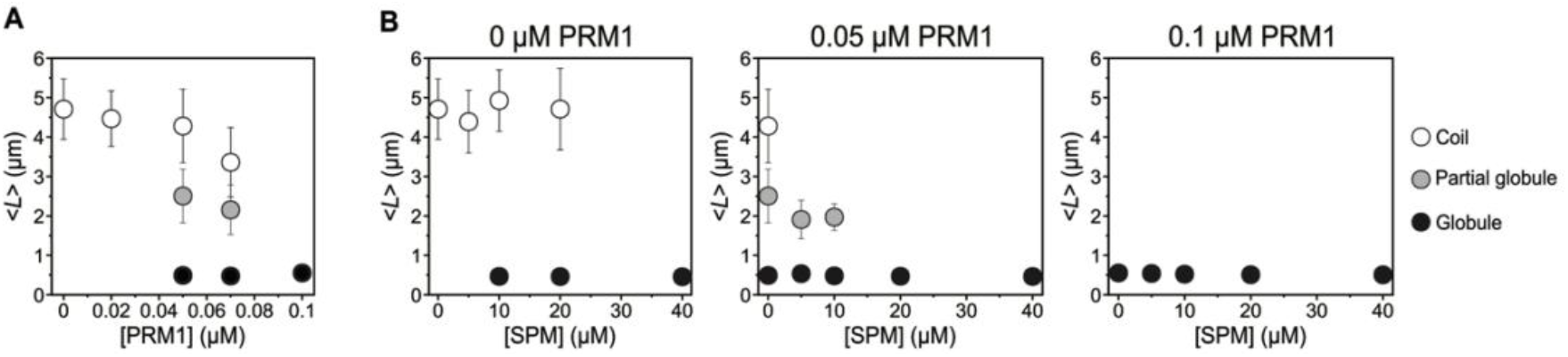
Ensemble-averaged long-axis length, <*L*>, of T4 GT7 DNA in each conformational state. Ensemble-averaged long-axis length, <*L*>, of coil, partial globule, and globule conformations as a function of PRM1 concentration (A) and as a function of SPM concentration (B) in the absence of PRM1 and in the presence of 0.05 and 0.10 µM PRM1, respectively. The distribution widths of *L* due to Brownian motion are represented by the standard deviation (SD). The three conformational states correspond to the white, gray, and black histograms shown in Fig. 1B.

## 4. Discussion

The present study demonstrates that human PRM1 and SPM induce distinct yet complementary modes of giant DNA compaction. Although PRM1 alone induced a continuous conformational transition through intermediate conformations, SPM alone induced a discrete coil–globule transition. Under coexistence conditions, however, SPM promoted structural progression along the PRM1-induced folding pathway, indicating complementary rather than independent roles of these two biologically relevant DNA-condensing molecules.

The complementary behavior observed here can be understood in terms of the distinct physicochemical mechanisms by which PRM1 and SPM compact DNA. Previous single-molecule studies demonstrated that protamines induce a continuous conformational transition through intermediate conformations, whereas small multivalent polyamines induce a discrete coil–globule transition [12, 13, 15]. The discrete folding transition induced by multivalent polyamines has been attributed to extensive charge neutralization of DNA by multivalent counterions, accompanied by counterion exchange and the resulting attractive interactions between DNA segments [12, 16]. In contrast, gradual DNA compaction induced by macromolecular polycations has been interpreted in terms of progressive changes in effective segment–segment interactions accompanying polycation binding. In the case of polyarginine, a theoretical treatment reproduced the continuous decrease in DNA size by assuming that increasing polycation binding progressively enhances attractive interactions between DNA segments [25]. Recent single-molecule and molecular simulation studies, primarily using salmon protamine, further indicate that protamine–DNA compaction involves multiple microscopic interaction modes and structures, including DNA bending, looping, flowers, loop stacks, and mechanically stable tangling [5, 23, 24, 26]. Thus, the gradual change in global DNA dimensions observed here does not necessarily represent a single microscopic folding mechanism, but may reflect the progressive population of multiple protamine-mediated DNA conformations.

The present observations extend these previous findings by demonstrating how these two distinct modes of DNA compaction complement one another under coexistence conditions. At 0.05 μM PRM1, SPM did not simply reproduce the discrete transition observed in the absence of PRM1. Instead, the disappearance of the coil population together with the progressive decrease in the partial globule population and concomitant increase in the globule population suggests that SPM promotes structural progression along the PRM1-induced folding pathway. The quantitative analysis of the long-axis length further supports this interpretation. Increasing PRM1 progressively shortened the coil and partial globule conformations, whereas SPM produced little change in the size of each conformational state after classification. Together, these findings indicate that PRM1 establishes a continuous folding pathway, whereas SPM promotes structural progression along this pathway toward the fully compact globule state. One possible interpretation is that progressive changes in effective segment–segment interactions induced by PRM1 alter the physicochemical landscape governing DNA compaction, such that SPM no longer induces its characteristic discrete coil–globule transition. Instead, SPM may enhance attractive interactions within the PRM1-generated conformational ensemble, thereby promoting structural progression along the continuous folding pathway toward the fully compact globule state. Further studies will be required to clarify the molecular basis of this cooperative mechanism.

During sperm maturation, histones are progressively replaced by protamines, leading to extensive chromatin condensation [1]. Polyamines have also been implicated in spermatogenesis and testicular physiology, where they are thought to contribute to chromatin organization and sperm function [27]. Recent studies have shown that protamine-mediated DNA interactions are strongly influenced by coexisting chromatin-associated proteins. In a system using salmon protamine, HMGB1 and transition protein 1 counteract the formation of mechanically stable protamine–DNA tangles and promote more dynamic protamine–DNA condensates [26]. In contrast, the present results demonstrate that SPM cooperates with PRM1 to promote progression toward compact globule conformations. Together, these observations suggest that protamine-mediated DNA compaction can be modulated in opposing directions by different components of the surrounding molecular environment. The present study employed naked giant DNA under simplified in vitro conditions and therefore does not directly reproduce the molecular environment of spermatogenic chromatin. Nevertheless, direct visualization of individual giant DNA molecules provides a platform for systematically examining how coexisting molecular and ionic factors alter protamine-mediated higher-order structural transitions. The present findings identify SPM as one such factor capable of promoting PRM1-mediated DNA compaction and provide a physicochemical basis for further investigating the multicomponent regulation of genome compaction during sperm maturation.

## 5. Conclusions

In conclusion, the present study demonstrates that human PRM1 and SPM play distinct yet complementary roles in giant DNA compaction. Human PRM1 induced a continuous conformational transition through intermediate conformations, whereas SPM induced a discrete coil–globule transition. Under coexistence conditions, SPM promoted structural progression along the PRM1-induced continuous folding pathway toward the fully compact globule state. These findings provide a physicochemical framework for understanding the distinct yet complementary roles of human PRM1 and SPM in higher-order structural transitions of giant DNA.

## Acknowledgements

This work was partially supported by JSPS KAKENHI Grant Numbers 23K14159 and 26K17112 to T. N.

## Author contributions

Takashi Nishio: Conceptualization, Methodology, Investigation, Formal analysis, Visualization, Writing – original draft, Writing – review & editing, Funding acquisition.

Yoshio Kato: Methodology, Resources, Writing – review & editing.

Both authors discussed the results and approved the final manuscript.

## Data availability

The data supporting this article are included in the main article and its Supplementary Information. Additional data are available from the corresponding author upon reasonable request.

