## Supplementary Information for "Complementary roles of human protamine 1 and spermine in giant DNA compaction"

**A**

Human protamine 1 (PRM1)

ARYRCCRSQSRSRYYRQRQRSRRRRRRSCQTRRRAMRCCRPRYRPRCRRH

**B**

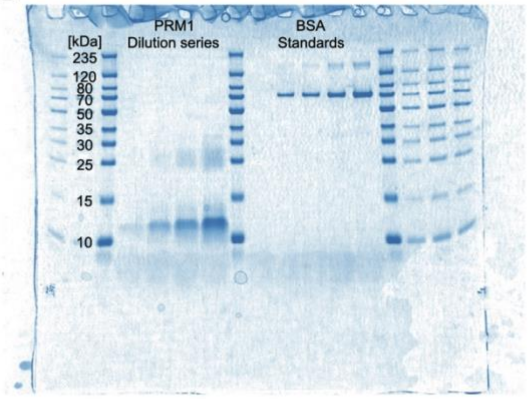


**Fig. S1. Recombinant human PRM1 used in this study.**

(A) Amino acid sequence of recombinant human PRM1 after removal of the N-terminal fusion tag. (B) Representative SDS-PAGE used for protein quantification. PRM1 samples were loaded at 0.1×, 0.25×, 0.5×, and 1× relative to the stock solution. BSA standards were loaded at 0, 0.0625, 0.125, 0.25, and 0.5 µg per lane. The concentration of the undiluted PRM1 stock solution (1×) was estimated to be 107.6 ng/µL (16.1 µM).
